# Mycobiome of *Pinus pinaster* trees naturally infected by the pinewood nematode *Bursaphelenchus xylophilus*

**DOI:** 10.1101/2025.02.21.639425

**Authors:** Cláudia S. L. Vicente, Ana Rita Varela, Anna Vetrainno, Margarida Espada, Maria de Lurdes Inácio

## Abstract

Fungi are important biological elements in the Pine wilt disease (PWD) complex. In the late stages of the disease, the pinewood nematode (PWN) *Bursaphelenchus xylophilus* feeds on the fungal flora available in the pine tree for survival and multiplication. Previous studies have confirmed a close relation between the PWN and blue-stain fungi (Ophiostomatales), which are necrotrophic pathogens associated with bark beetles (Coleoptera: Scolytidae). The PWN is able to grow densely in the presence of these fungi, which results in a higher number of nematodes transferred to the insect-vector *Monochamus* spp. To understand the spatial diversity and structure of *Pinus pinaster* mycobiome, wood samples from PWN-infected and non-infected pine trees were collected in three locations of Continental mainland Portugal with different PWD records, during the maturation phase of the insect-vector *M. galloprovincialis* (winter 2019-spring 2020). The PWN-mycobiome from the PWN-infected *P. pinaster* was also characterized. A total of 27 samples of *P. pinaster* and 13 samples of PWN from PWN-infected trees were characterized using ITS2 amplicon sequencing. The diversity and structure of the fungal communities in *P. pinaster* varied with disease status suggesting that the PWN presence affects the endophytic fungal communities. For both *P. pinaster* and PWN fungal communities, differences were also associated with locations (recent PWD loci Seia, and long-term PWN locus Companhia das Lezírias and Tróia). Ophiostomatales were mainly detected in PWN-infected *P. pinaster*. This research contributes to increase the knowledge on the ecology of the fungal communities in PWD complex.

## Introduction

Pine wilt disease (PWD) is caused by the plant-parasitic pinewood nematode *Bursaphelenchus xylophilus* (also known as pinewood nematode, PWN), which is originated from North America. Native American conifers are mostly resistant or tolerant to *B. xylophilus*, not causing significant damage, other than on exotic tree species [1]. In early 20^th^ century, PWN was introduced in Japan, spreading afterwards to other Asian countries such as China, Korea and Taiwan [1]. Many native tree species in this region, such as *Pinus thunbergii* (Japanese black pine) and *P. densiflora* (Japanese red pine), are highly susceptible to PWN and the damage caused is extensive [2]. In Europe, PWN was detected for the first time in 1999 in Portugal and later in Spain [3-6]. Currently, PWN is a European A2 quarantine organism listed as a top pathogen that causes economic and ecological damage to the coniferous forestry industry, including restrictions in the circulation of wood products [7]. The host range of susceptible species to PWN also includes *P. pinaster, P. nigra, P. radiata* and *P. sylvestris* [8, 11]. PWN is a necrotrophic plant-parasitic nematode with a complex life cycle which includes the ability to feed on living plant cells (phytophagous phase) and tree endophytic fungi (obligatory mycetophagous phase) [2,12-15]. This nematode established a specific phoretic relation with the insect-vector *Monochamus* spp., mostly involved in its dispersion from dead to healthy pine trees [2]. The nematode migrates, feeds and reproduces inside of the plant living tissues, which results in the tree’s physical destruction [2, 16]. In an early stage of the infection, the nematode feeds on the parenchyma cells of the ray canals and resin ducts in the cortex and in the xylem. At a later stage, the nematode population increases being detected in the cortex, phloem, cambium, and xylem tissues. At this point, the nematode causes the disruption of the water flow, blocking of the vascular system (embolism) and cavitation leading to wilting symptoms and tree death [2, 17-18]. Once the host tree starts declining, the nematode switches to its fungal feeding strategy (mycetophagous phase) [14, 15]. Several studies have shown that the favorable food for PWN are beetle-associated fungi belonging to order Ophiostomatales [17] (further detailed in 19). Some of these ophiostomatoids, crucial for PWN multiplication inside the host tree and in the insect-vector, are also known as blue-stain fungi and are responsible for the blue coloration of the weakened pine trees. The presence of blue-stain enhances nematode reproduction rates, indicating a probable commensal relationship between the nematode and specific fungal species [20]. Just before the departure of the insect-vector from the dying tree, the PWN enters in a dispersal mode, referred to as *dauer* stage, that enables their survival during tree-to-tree transmission [2].

The concept of tree holobiont has been modernized by the availability of high-throughput sequencing technologies (HTS), such as metagenomics, that have allowed a holistic point-of-view into the complexity of microbial interactions with tree host [21-22]. In the context of PWD, the impact of the PWN in the microbial (bacteria/fungi) communities have been addressed on several *Pinus* species, emphasizing the cascade effect of nematode infection in the diversity and composition of these communities [23-24]. In the current study, we investigate the structure of the mycobiome mycobiome non-infected and PWN-infected *Pinus pinaster* trees in three biogeographical locations in mainland Portugal (Companhia das Lezírias; Tróia and Seia). We hypothesize that: i) fungal communities of *P. pinaster* shift in diversity and composition in response to PWN infection; ii) fungal communities of PWN share a similar composition and structure to those of their host tree; and iii) the sampling locations may play a driving factor in shaping the *P. pinaster* mycobiome. This is the first study to focus solely in the mycobiome of *P. pinaster* trees and their plant-parasitic nematode *B. xylophilus*, highlighting the variation and vulnerability of fungal communities and ecosystem functions in the presence of a pathogenic agent.

## Results

### Sequencing data and diversity estimates

From a total of 30 pine samples, only 27 were considered for further analysis due to low number of reads. From these, a total of 1,605,754 high-quality reads were obtained in the three sampling sites, after denoising and quality filtering. From the PWN dataset, a total of 13 PWN samples were analyzed, resulting in 778,959 high-quality reads. In both datasets, the sequencing depth was sufficient to describe the diversity and richness of fungal communities as indicated by the rarefaction curves’ plateau (Supplemental Figure S1a and S1b). The estimated diversity indexes for both datasets are presented in Table 1 and 2.

**Table 1.** Alpha-diversity estimations for fungal communities from *Pinus pinaster* samples. Statistical analysis was performed between locations (C. Lezírias; Seia and Tróia) within each condition (“PWN-infected”; “non-infected”). Different superscript letters (a/b) indicate significant differences at p<0.05 (estimated with Kruskal-Wallis test). Data are means ± SE (standard error).

| Condition | Location | Observed ASVs |  | Chao1<sup>1</sup> |  | Simpson<sup>2</sup> index |  | Shannon<sup>2</sup> index |  | Pielou's index |  | GD<sup>2</sup> |  |
| --- | --- | --- | --- | --- | --- | --- | --- | --- | --- | --- | --- | --- | --- |
| PWN-infected | C. Lezírias | 30.4(3.4) | b | 31.0(4.0) | b | 0.70(0.0) | a | 1.71(0.13) | b | 0.49(0.0) | a | 1 | a |
|  | Seia | 34.6(4.7) | b | 35.4(4.9) | b | 0.83(0.0) | a | 2.26(0.14) | b | 0.65(0.0) | b | 1 | a |
|  | Tróia | 86.3(16.9) | a | 95.8(22.7) | a | 0.82(0.0) | a | 2.69(0.30) | a | 0.59(0.0) | b | 1 | a |
| Non-infected | C. Lezírias | 176.0(27.9) | a | 180.0(29.5) | a | 0.96(0.0) | a | 4.18(0.13) | a | 0.82(0.0) | a | 1 | a |
|  | Seia | 132.0(29.1) | a | 141.0(29.1) | a | 0.80(0.1) | a | 2.61(0.40) | b | 0.53(0.1) | b | 1 | a |
|  | Tróia | 190.0(31.8) | a | 203.0(32.6) | a | 0.89(0.1) | a | 3.51(0.39) | b | 0.66(0.1) | b | 1 | a |

<sup>1</sup>Species level: 99% similarity threshold to define average sequence variants (ASVs); Chao1: Chao's species richness estimator;
<sup>2</sup>Simpson and Shannon index indicate, respectively, Simpson diversity index (1-D) and Shannon diversity index; GD, Good's coverage.

**Table 2.** Alpha-diversity estimations for fungal communities of *Bursaphelenchus xylophilus* samples. Statistical analysis was performed between locations (C. Lezírias; Seia and Tróia). Different superscript letters (a/b) indicate significant differences at p<0.05 (estimated with Kruskal-Wallis test). Data are means ± SE (standard error).

| Location | Observed ASVs (SE) |  | Chao1<sup>1</sup> (SE) |  | Simpson index |  | Shannon<sup>2</sup> index |  | Pielou index |  | GD<sup>2</sup> |  |
| --- | --- | --- | --- | --- | --- | --- | --- | --- | --- | --- | --- | --- |
| C. Lezírias | 140(18.3) | a | 145(19.6) | a | 0.88(0.1) | a | 3.34(0.3) | a | 0.67(0.1) | a | 1 | a |
| Seia | 72.5(6.5) | b | 73.8(7.1) | b | 0.81(0.0) | a | 2.38(0.2) | a | 0.56(0.0) | a | 1 | a |
| Tróia | 84.5(21.9) | b | 89.8(22.9) | b | 0.55(0.1) | b | 1.54(0.5) | b | 0.35(0.1) | b | 1 | a |

<sup>1</sup>Species level: 99% similarity threshold to define average sequence variants (ASVs); Chao1: Chao's species richness estimator;
<sup>2</sup>Simpson and Shannon index indicate, respectively, Simpson diversity index (1-D) and Shannon diversity index; GD, Good's coverage index.

Overall, samples from both datasets presented a Good’s coverage index (0.99-1.00), corroborating the results from the rarefaction curve analysis (Supplemental Figure S1). For all collection sites in the *P. pinaster* dataset, samples from “PWN-infected” trees presented a lower number of ASVs (C. Lezírias, 22-42 ASVs; Seia, 18-45 ASVs; and Tróia, 59-162 ASVs), in contrast with “non-infected” trees that were richer with more ASVs per sample (C. Lezírias, 124-249 ASV; Seia, 74-206 ASV; and Tróia, 80-256) (Table 1 and Supplemental Table S1). Likewise, fungal communities from “PWN-infected” trees presented lower diversity (0.70-0.83) than “non-infected” ones, as indicated by the Simpson’s and Shannon’s diversity indexes. For “PWN-infected” condition, statistical differences (p<0.05) were denoted between locations (C. Lezírias, Seia, and Tróia) in almost all indexes, with exception of Simpson’s (Table 1). Quite the opposite, for the “non-infected” condition, only Shannon and Pielou’s indexes were significant different (p<0.05) between C. Lezírias and the other two locations (Table 1). The richness of the fungal communities associated with *B. xylophilus’* (Table 2), assessed by the number of ASVs was statistically higher (p<0.05) in C. Lezírias (140±18.3) than in Seia (72.5±6.5) and Tróia (84.5±21.9). However, in terms of diversity, the results followed an opposite trend, with fungal communities from C. Lezírias and Seia being more similar to each other than those from Tróia.

Following our previous results based on cultural-dependent isolation of fungal communities [24], we hypothesized that “PWN-infected” and “non-infected” mycobiome differ not only due to the presence of the PWN and disease progression but also as a result of the variations in biogeographic patterns aligned with the timing of disease introduction in the different collection sites. To test these hypothesis, beta-diversity was statistically analyzed (PERMANOVA) (Table 3).

**Table 3.** perMANOVA statistical tests on beta-diversity metrices (Bray-Curtis distance). Two metadata categories were considered, namely conditions (“PWN-infected” and “non-infected”) and Location (C. Lezírias, Seia and Tróia).

| Dataset | Metadata Category | perMANOVA |  |
| --- | --- | --- | --- |
|  |  | Pseudo-F | p-value |
| *Pinus pinaster* | Condition | 4.434 | **0.001** *** |
|  | Location | 2.309 | **0.001** *** |
| *Bursaphelenchus xylophilus* | Location | 4.199 | **0.001** *** |

\*Asterisks indicate significant differences: $p < 0.001$ , \*\*\*; $p < 0.01$ , \*\*; $p < 0.05$ , \*.

Our premise that fungal communities difference correlate with the presence of PWN (p=0.001), as well as with the collection site (p=0.001) were corroborated (Table 3). For the PWN-dataset, statistical differences were observed between collection sites (PERMANOVA, Pseudo-F = 4.199, p=0.001).

### Mycobiome composition, abundance and structure of *Pinus pinaster*

The composition and abundance of the fungal communities of *Pinus pinaster* is presented in Figure 1. Within Fungi kingdom, sequences were assigned onto 1067 ASVs within Fungi kingdom, compromising 5 phyla, 77 orders and 218 genera. Overall, only 141 ASVs were shared between “PWN-infected” and “non-infected” trees, regardless the sampling location (Fig. 1a). Supporting the results obtained from the diversity and richness estimators, the “non-infected” trees presented unique 775 ASVs, while “PWN-infected” trees presented only 151 ASVs (Fig. 1a). The fungal composition of *P. pinaster* differed between “PWN-infected” and “non-infected” trees, with communities from C. Lezírias and Seia being more similar to each other than to those from Tróia (Fig.1b). The most abundant phyla, in both “non-infected” and “PWN-infected” trees, were Ascomycota (56.2-82.4%) and Basidiomycota (16.4-43.2%). In “PWN-infected” trees, for both Seia and Lezírias, the most common orders for were Saccharomycetales (Ascomycota, 54.4-62.7%), followed by Ophiostomatales (Ascomycota, 12.9-16.0%) and Atratiellales (Basidiomycota, 13-14,4%) (Fig. 1b). In Tróia, the most predominant orders were Hymenochaetales (Basidiomycota, 21.8%), Coniochaetales (Ascomycota, 15.0%), Saccharomycetales (Ascomycota, 11.4%) and Ophiostomatales (Ascomycota, 3.63%). The most abundant genera, in Saccharomycetales order, was *Nakazawaea* (1.46% in Tróia; 29.5% in Seia; and 30% in C. Lezírias), while in Ophiostomatales order, the most common genera were *Ophiostoma (*0.83% in Tróia; 5.65% in Seia; and 2.91% in C. Lezírias), *Leptographium* (0.28% in Tróia; 2.37% in Seia; and 6.37% in C. Lezírias); and *Ceratocystiopsis* (0.90% in Tróia; 3.12% in Seia; and 4.59% in C. Lezírias). In “non-infected” trees, the most abundant orders were for Tróia, unclassified phylum Ascomycota (un_p_Ascomycota; 24.5%), Capnodiales (14.3%) and Hymenochaetales (11.9%); for Seia, Pleosporales (38.5%), Russulales (14.9%) and Helotiales (12.1%); and for C. Lezírias, unclassified phylum Ascomycota (29.4%), Capnodiales (21.9%), and Tremellales (8%). The orders Saccharomycetales and Ophiostomatales were not detected or less than 5% prevalent.

**Figure 1.**
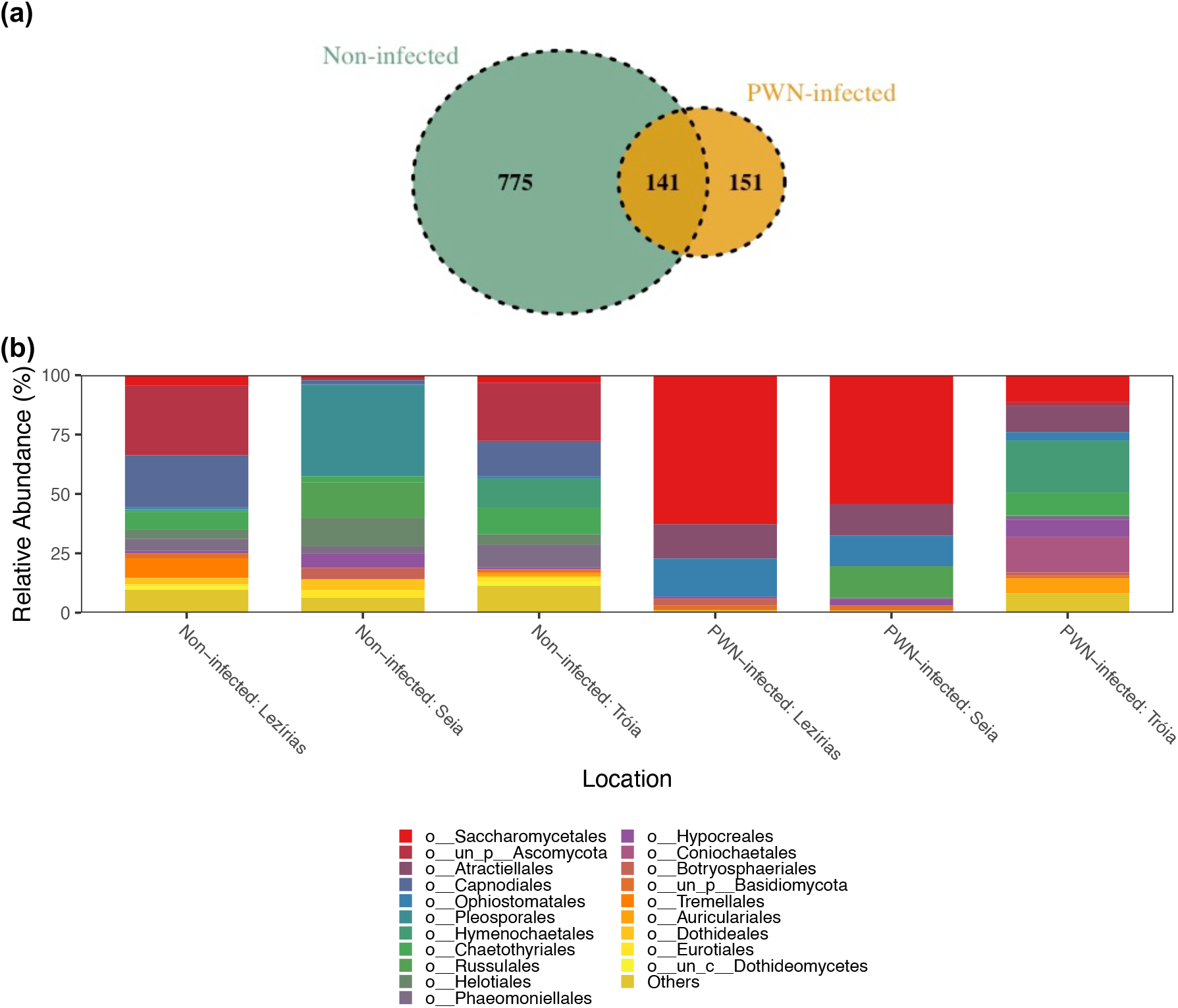
Composition and abundance of fungal communities of *Pinus pinaster*: a) Venn diagram representing the distribution of ASVs across conditions (“non-infected” and “PWN-infected” trees); and b) relative abundance of the top 20 orders for each condition (“non-infected” and “PWN-infected” trees) and location (C. Lezírias, Seia and Tróia).

A principal co-ordinates analysis (PCoA) was performed to ascertain fungal communities’ structure in terms of sampling location and the presence of the PWN (Fig. 2). The first axis (PCoA 1) explains 21.33% of the variation between samples and clearly separates fungal communities of “non-infected” trees from the “PWN-infected” trees. The second axis (PCoA2) only explains 12.6% of the variation in the distribution. In fact, among “non-infected” trees, fungal communities tend to cluster by sampling location, being ASV0021 and ASV0293 (both Pleosporales order, Didymellaceae family, unidentified genus) related with the distribution of Seia while ASV0626 (Capnodiales order) related with Tróia and C. Lezírias. Among “PWN-infected” trees, all samples tend to group regardless the sampling location, being the ASV0014 (genus *Proceropycnis*) and ASV0131 (genus *Nakazawaea*) related with their distribution. Together, these axes explain 33.93% of the variability observed between conditions and locations.

**Figure 2.**
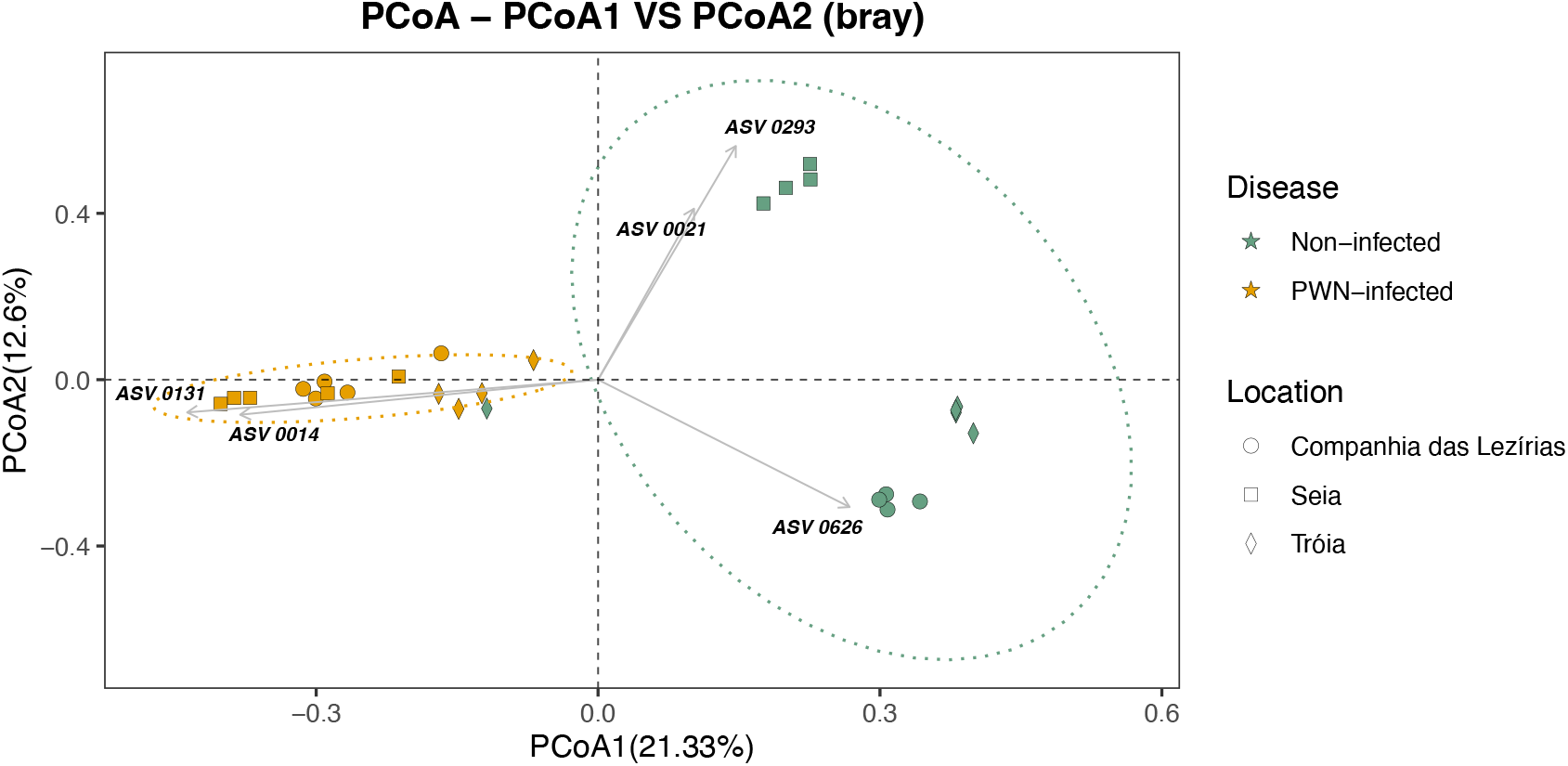
Principal coordinates analysis (PCoA) of fungal communities of *Pinus pinaster* using Bray-Curtis distance matrix. The followings ASVs are related with communities’ distribution: ASV0014 (Basidiomycota, Atractiellomycetes, Atractiellales, Hoehnelomycetaceae, *Proceropycnis*); ASV0021 (Ascomycota, Dothideomycetes, Pleosporales, Didymellaceae); ASV0131 (Ascomycota, Saccharomycetes, Saccharomycetales, Pichiaceae, *Nakazawaea*); ASV0293 (Ascomycota, Dothideomycetes, Pleosporales, Didymellaceae); ASV0626 (Ascomycota, Dothideomycetes, Capnodiales).

To validate previous results, a SIMPER analysis was performed to identify the ASVs that represented each condition and location (Supplemental Table S2). Considering only these two conditions, the average similarity within PWN-infected trees was 20.06%, while within “non-infected” trees was 18.96%. These results reflect the heterogeneity of the locations and the trees sampled. More importantly, the average dissimilarity between-condition was 94.59% that emphasizes the differences between PWN-infected and non-infected communities. The top ASVs contributors for this dissimilarity were ASV0014, ASV0131, also assigned in the PCoA analysis, and ASV0502 (Ascomycota, Saccharomycetes, Saccharomycetales, Pichiaceae, *Nakazawaea*, unidentified species) (Supplemental Table S2).

### Occurrence of PWN in the sampled trees

The detection of PWN for each tree sampled was determined by 48h-extraction from wood pellets, followed by nematode counting [24]. Intriguing, using ITS2 metagenomics, the fungal community of sample T9 from “non-infected” trees of Tróia showed a composition/abundance closer to “PWN-infected” trees and grouped within this condition, according with the PCoA analysis. To validate PWN classes previously considered, total DNA from each pine sample sequenced was also used for qPCR with a specific probe for *B. xylophilus* detection (Table 4). Overall, the qPCR results were in accordance with previous detection. For most of the samples, the higher PWN classes (e.g., higher number of PWN extracted) resulted on lower CT values, indicating early detection in qPCR amplification curve. However, the CT value for sample T9 (19.44) was unexpected since PWN class was 0 (e.g., no nematodes extracted), suggesting a positive detection of PWN. Thus, clustering of T9 within the “PWN-infected” trees was supported.

**Table 4.**
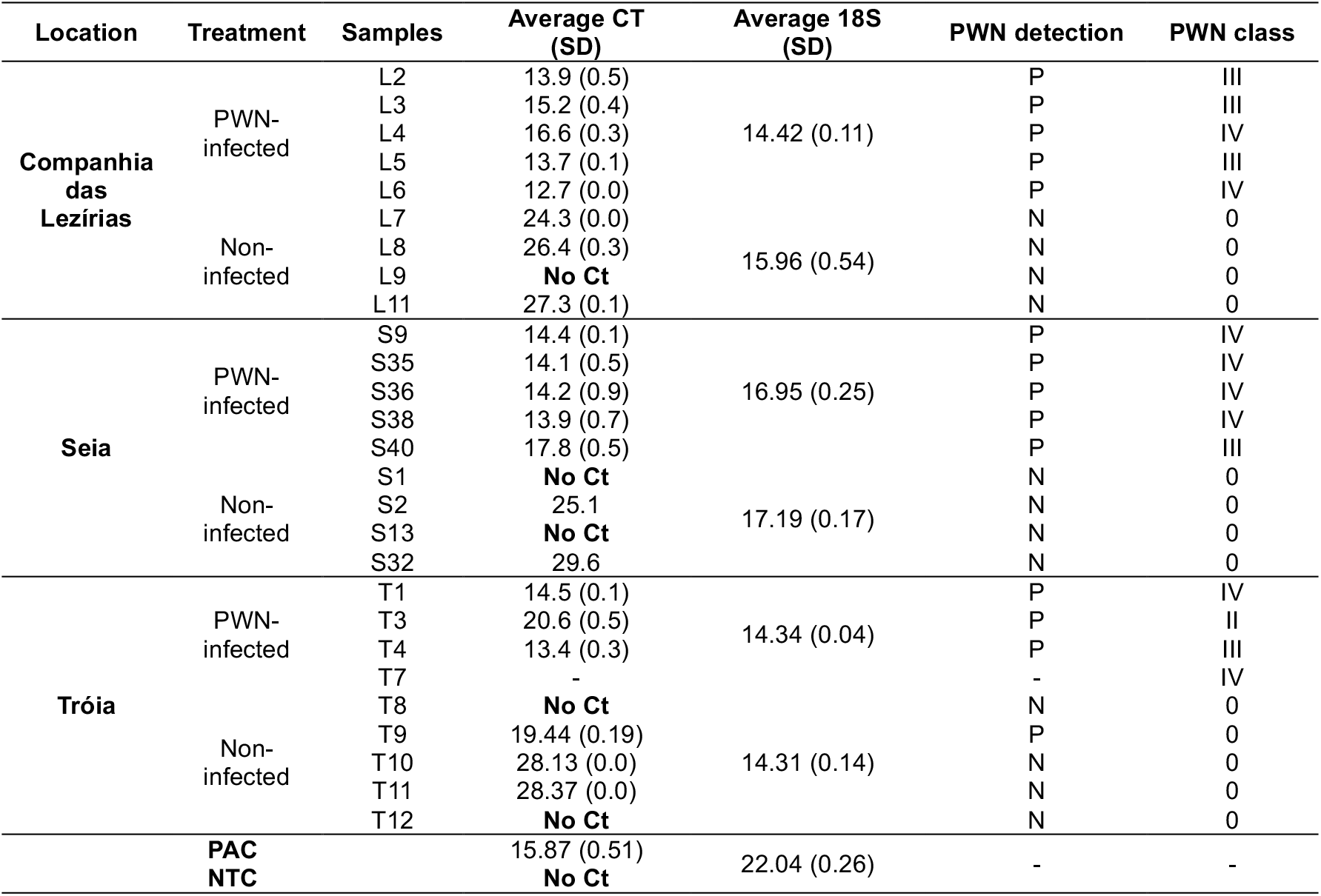
Detection of *Bursaphelenchus xylophilus* in wood samples by real-time PCR. The cut-off values recommended for the detection of *B. xylophilus* on wood extracts using real-time PCR from PM 7/4 (4) *Bursaphelenchus xylophilus* [7]. Positive amplification control (PAC) and negative template control (NTC) were considered to assign positive (P) or negative (N) PWN detection. PWN classes were previously determined by [22].

### Mycobiome communities’ composition, abundance and structure of *Bursaphelenchus xylophilus*

From PWN dataset, all sequences were assigned onto 616 ASVs and classified in 5 phyla, 57 orders, and 147 genera. From these, only 58 ASVs were shared between all three locations (Fig. 3a). The main phyla describing the fungal communities of *B. xylophilus* are Ascomycota (56.18-95.69%) and Basidiomycota (2.31-39.81%). The most predominant orders were different according to sampling location: in Tróia, Orbiliales (Ascomycota, 51.1%), Pezizomycotina_ord_Incertae_sedis (Ascomycota, 27.97%) and Saccharomycetales (Ascomycota, 12.01%); in Seia, Sporidiobolales (Basidiomycota, 34.38%), Pleosporales (Ascomycota, 25.1%), Saccharomycetales (Ascomycota, 15,34%); and, in Lezírias, Saccharomycetales (Ascomycota, 24.11%) Spizellomycetales (Chytridiomycota, 12.3%) and Capnodiales (Ascomycota, 9.31%) (Fig. 3a). The order Ophiostomatales was also detected in all samples but its abundance was less than 1%. On genus level, the most abundant and identified genera, according to sampling location, were: in Tróia, *Ciliophora* (Ascomycota, 28.0%) and *Arthrobotrys* (Ascomycota, 15.6%); in Seia, Rhodosporidiobolus (Basidiomycota, 30.9%), *Alternaria* (Ascomycota, 6.6%) and Groenewaldozyma (Ascomycota, 5.7%); and, in Lezírias, *Cladosporium* (Ascomycota, 7.7%), *Nakazawaea* (Ascomycota, 6.6%) and *Fusarium* (Ascomycota, 4.6%).

**Figure 3.**
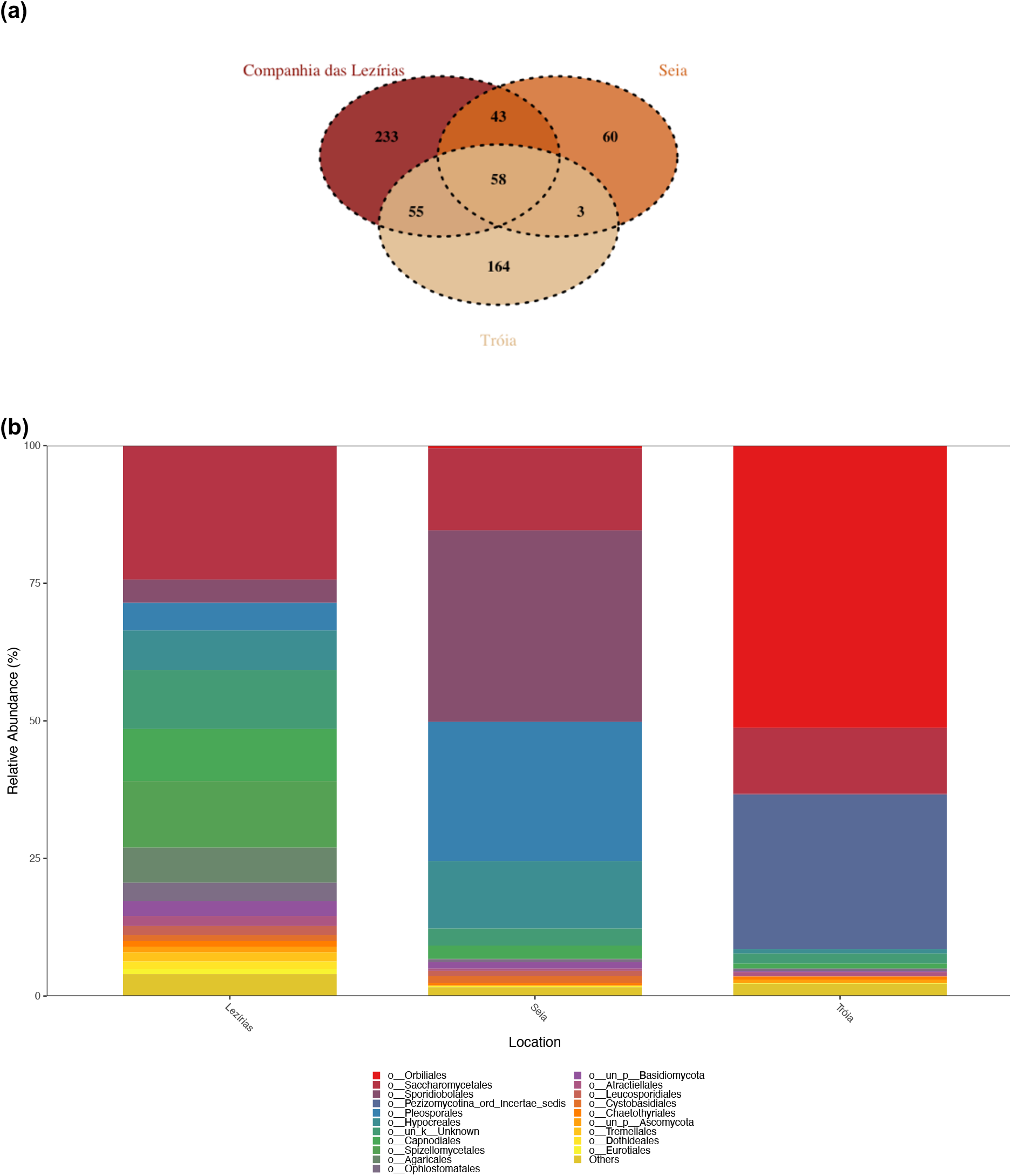
Composition and abundance of fungal communities of *Bursaphelenchus xylophilus*: a) Venn diagram representing the distribution of ASVs across conditions; and b) relative abundance of the top 20 orders for each location (Lezírias, Seia and Tróia).

The PCoA analysis for the *B. xylophilus* fungal communities is presented in Fig. 4. The first axis (PCoA1) explains 32.29% of the variation and clearly separates Tróia communities from Seia and Lezírias, while the second axis, with 14.92% of the variation, separates communities from Lezírias and Seia. The spatial distribution of Tróia’s fungal communities was driven by the ASV0140 (Orbiliales order Orbiliaceae family), the ASV0066 (genus *Ciliophora*) and ASV0458 (*Arthrobotrys cladodes*). Fungal communities from Seia were influenced by the abundance of ASV0543 (*Pseudopithomyces* genus) and ASV0606 (Pleosporales order, Didymellaceae family). Both axes explain 47.21% of the variability observed between locations.

**Figure 4.**
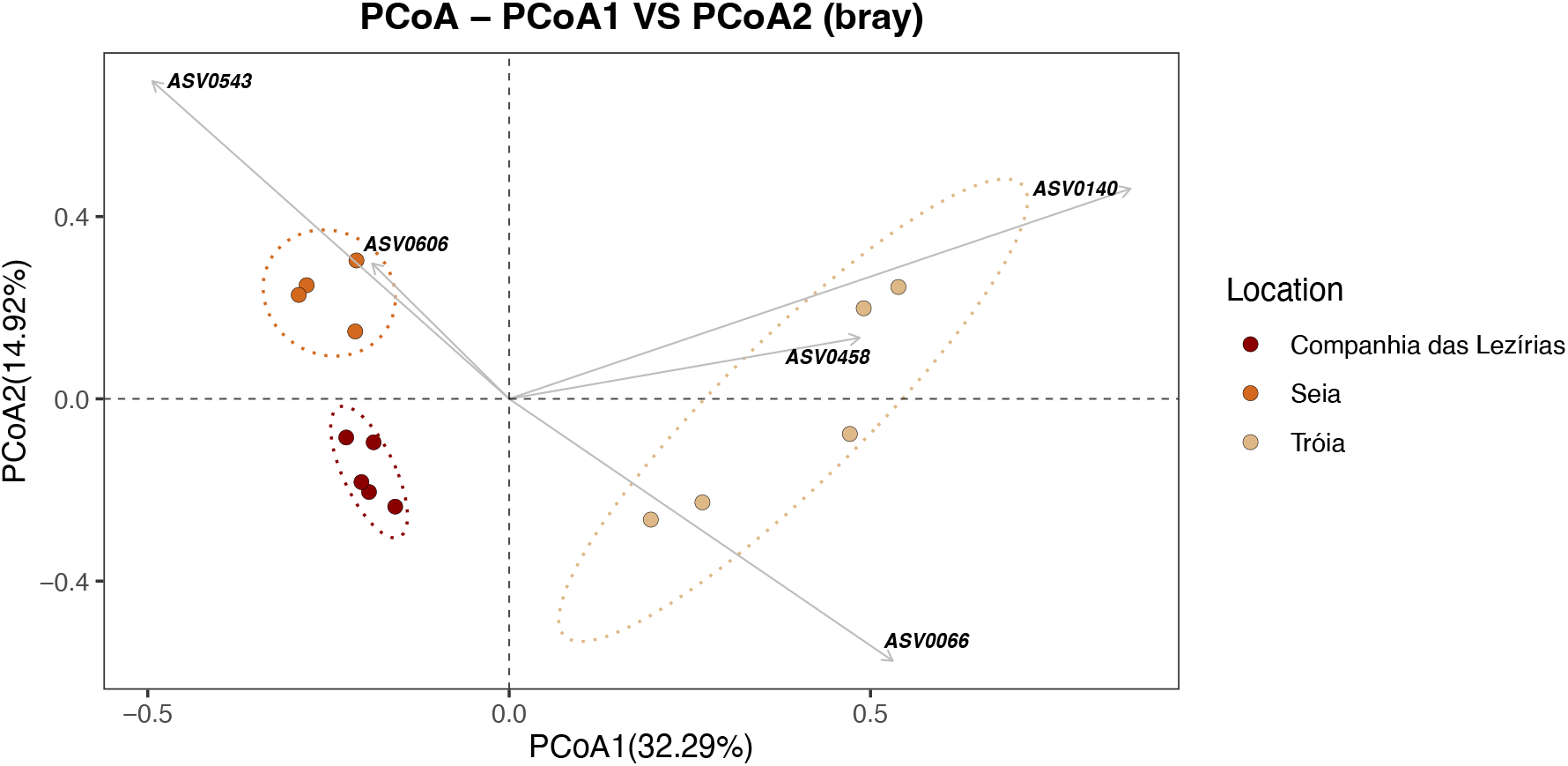
Principal coordinates analysis (PCoA) of fungal communities of *Bursaphelenchus xylophilus* using Bray-Curtis distance matrix. The followings ASVs are related with communities distribution: ASV0066 (Ascomycota, Pezizomycotina_cls_Incertae_sedis, Pezizomycotina_ord_Incertae_sedis, Pezizomycotina_fam_Incertae_sedis, *Ciliophora*); ASV0140 (Ascomycota, Orbiliomycetes, Orbiliales, Orbiliaceae); ASV0458 (Ascomycota, Orbiliomycetes, Orbiliales, Orbiliaceae, *Arthrobotrys, Arthrobotrys cladodes*); ASV0543 (Ascomycota, Dothideomycetes, Pleosporales, Didymosphaeriaceae, *Pseudopithomyces*); and ASV0606 (Ascomycota, Dothideomycetes, Pleosporales, Didymellaceae).

As seen for *P. pinaster* dataset, the SIMPER analysis is in accordance with the PCoA analysis for the *B. xylophilus* dataset (Supplemental Table S3). The average similarity within-locations ranged between 29.96% (Tróia) and 43.51% (Seia). The average dissimilarity between-locations were considerably high (71.08-89.46%), highlighting their distribution in the Fig. 4. For instances, ASV0543 and ASV0606 were among the most representative to separate fungal communities of Lezírias and Seia (as in the PCoA). In the case of the *B. xylophilus’* fungal communities of Lezírias and Tróia, the ASV0140, ASV0066 and ASV0458 were the highest contributors, which are also assigned in Fig. 4.

## Discussion

PWD is often explained using a disease triangle framework, which implicates the interaction of the host tree, insect-vector, and causal agent (PWN) as the main drivers of the disease. Recent research on plant disease dynamics suggests that the disease triangle model is transitioning into a pyramid-shaped, with the disease outcome being the result of the interactions between the host, host microbiome, pathogen, and environment [25]. Considering the transition of this paradigm, in this study, we analyzed the mycobiome of PWN-infected and non-infected *P. pinaster*, and also the mycobiome of the PWN, in three distinct locations in Portugal. Supporting our previous results on culturable fungal communities of PWN-infected *P. pinaster* [24], this work showed that fungal communities from both PWN-infected and non-infected maritime pines were significantly different in terms of species richness, diversity and composition, and that these differences were observed among sampling sites (contrasting biogeographical locations). The fungal communities of the PWN among the three locations were also significant diverse and resembled, as expected, the communities of their host trees. Thus, the presence of the PWN caused a significant shift in: i) the native mycobiome of *P. pinaster*, as seen in other pine species [26-33] or in other microorganisms’ communities [23]); and ii) in the abundance of Ophiostomatales, almost exclusively in PWN-infected pine trees, emphasizing the close relation between these organisms [19, 27-33] The observed interaction between PWN and these blue-stain fungi suggests a long-term co-evolutionary relationship.

Host tree, geography and climate were predicted as the main drivers of insect and fungal communities’ structure in 155 tree species in 32 countries (6 continents) [34]. In the context of PWD, Liu et al. [26] characterized the fungal communities of soil, root, and needles of *P. thunbergii* infected and non-infected with PWN, showing that nematode infection only affected needle fungal communities, with prevalence of Ascomycota phylum, genera *Diplodia* and *Hormonema*. In *P. massoniana*, significant differences in fungal abundances between healthy and wilted pine trees were also reported in different geographical locations in Zhejiang Province (China) [27-28]. Specific genera from wilted trees were reported such as *Cryptoporus* (Basidiomycota, Polyporales), *Graphilbum* and *Sporothrix* (both Ascomycota, Ophiostomatales) and *Geosmithia* (Ascomycota, Hypocrales), that were also seen in resistant *P. caribaea, P. elliottii*, and *P. taeda* [28]. Ophiostomatales fungi were only found in wilted *P. massoniana*, specifically in the *Graphilbum* genus [27]. The phyllosphere fungal communities of *P. tabuliformis* were also affected by PWN infection at early and late stage of the disease, represented by *Septoria, Devriesia, Truncatella* and *Pestalotiopsis* [31]. More recently, Zhang and co-workers [32] explored the gallery microbiomes associated with *M. alternatus* concluding also that PWN infection altered the fungal communities assembles with major representation of Ophiostomataceae family.

Yeasts, unicellular fungus, play important roles as symbionts of bark beetles, helping them in food localization and digestion, in the detoxification of plant secondary metabolites, and even in the regulation of phoretic relations with other organisms [35]. In our study, yeasts from genus *Nakazawaea* (Ascomycota phylum) contributed for the dissimilarity between “non-infected” and “PWN-infected” *P. pinaster* communities. Yeasts belonging to this genus are generally described as saprophytes and were isolated from different sources such as the insect’s gut [36] or the phylloplane of sugar cane leaves [37]. Furthermore, the genus *Nakazawaea* is commonly associated with the gut microbiome of different species of bark beetles (Coleoptera: Curculionidae: Scolytinae) [38]. During the wilting process of an PWN-infected pine tree, bark beetles and pine sawyer beetles are chemically-attracted for the host tree around the same time. These bark beetles also transport several species of ophiostomatoid fungi attached on the exoskeleton surface or in specialized organs [19]. Not surprisingly, the mycobiome of these bark beetles is constituted by fungal assemblages similar to the PWN-infected *P. pinaster* here reported, in addition to being reported in the pupal chambers of *M. alternatus* or in the third-stage dispersal juveniles of *B. xylophilus* [33]. Our results are consistent with other observations and accentuate the PWN effect on the holobiont community of the pine tree, as well as the dispersion of other communities by related-insects.

Presently, HTS technologies are considered paramount for the molecular diagnosis of plant pathogens, not only for routine analysis application but also for the surveillance and prevention of newly potential introductions [39-40]. In this work, PWN was detected in the libraries generated for PWN-infected trees (data not shown), and in one “non-infected” tree (T9), which was further validated by real-time PCR [7]. The lack of detection of PWN though the conventional extraction method showing that no alive nematodes were present may indicate a case of aborted infection event, or, as reported elsewhere [41], genetic resistance of the individual *P. pinaster* tree that conferred resistance to nematode infection. In his study, Kovalchuck et al. [41] analyzed the mycobiome of asymptomatic and symptomatic Norway spruce trees infected by *Heterobasidion* spp., denoting that, although infected with the presence of the pathogenic agent, some individual trees did not develop the disease. The authors argued two possibilities for this result: sampling collection time-point in the initial stages of the disease development, or the intrinsic genetic resistance of individuals trees [41]. Either way, the use of HTS technologies help to tangle these experimental limitations in both studies.

Aside of the stochastic events, the complexity of the PWD and the cascade effects of the PWN on the tree holobiont urges the need to consider fungal communities, specifically Ophiostomatales and yeast, as important contributors for disease development. These fungi play a crucial role in shaping the intricate interactions within this ecological system, influencing nematode reproduction, the behavior of insect vectors, and overall disease dynamics. Understanding the function of fungi in the PWD complex is essential for effective disease management and control strategies. This study builds on the work initiated by Vicente et al. [24], pioneering research into the mycobiome of *P. pinaster* and the pinewood nematode at a European level. It marks a significant contribution to understanding the fungal communities associated with this host-pathogen system.

## Methods

### Sample collection and processing

PWD-symptomatic and asymptomatic *P. pinaster* trees were sampled, at mid-trunk, in three contrasting regions in mainland Portugal with different PWD records, namely, long-term PWD foci, Tróia (T) and Companhia das Lezírias (Ribatejo) and a more recent or transitional PWD foci, in Seia (S). The surveys were performed between Winter 2019 and Spring 2020, in-between the period of the insect-vector’ maturation feeding as referred in our previous work [24]. Wood samples were processed as described in [24]. After confirmation of PWN infection, PWD-symptomatic and asymptomatic *P. pinaster* trees were assigned to the PWN-infected and non-infected *P. pinaster* trees. Wood samples and nematode suspensions were flash-frozen with liquid nitrogen and stored at -80ºC until DNA extraction.

### Total DNA extraction, sequencing and quality control

Frozen wood samples were powdered using the Retsch Ultra Centrifugal mill ZM 200 (Retsch, Germany), properly cleaned with commercial sodium hypochlorite and sterilized with 70% EtOH (v/v), between samples to prevent cross-contamination. Total DNA extraction of wood powder was conducted using the modified cetyl-trimethyl ammonium bromide (CTAB)-based method described by Terhonen et al. [42]). In the case of nematode suspensions (pool of nematodes extracted from PWN-infected *P. pinaster*), and prior to DNA extraction, nematodes were pelleted and washed with sterilized distilled water 3x to remove wood debris. Total DNA extraction of PWNs was performed using DNeasy Blood & Tissue kit (QIAGEN, Valencia, CA, USA) according to the manufactures’ instructions. The concentrations and purity of the isolated DNA were measured by Qubit 4.0 fluorometer (Thermo Fisher Scientific, USA) and NanoDrop TM 2000 spectrophotometer (Thermo Fisher Scientific, USA), respectively. The presence of amplifiable DNA was confirmed by PCR amplification using the primers gITS7 and ITS4 [43]. A total of 45 DNA samples were selected and sent for sequencing on Illumina MiSeq 2x 300bp (Illumina, Inc. San Diego, CA, USA) at EUROFINS Genomics (Cologne, Germany). Raw ITS2 Illumina data was demultiplexed and quality filtered at QIIME2 (version 2022.11) [44] following the standard pipeline analysis. For both datasets (wood and nematode samples), pair-end reads were denoised using DADA2 plugin [45] and amplicon sequence variants (ASV) determined after discarding biased reads (e.g., chimeras).

### Taxonomic assignment, diversity and statistical analysis

Denoised ASVs were taxonomically assigned at 99% similarity using classify-sklearn plugin [46] against UNITE database (version 9.0) [47]. ASVs classified as Viriplantae, Metazoa, Alveolata, Heterolobosa, Rhizaria, Stramenopila, Protista and unassigned were filtered. After, ASVs table for both wood and nematode samples were rarefied, respectively, at 8000 and 15000 sequences per sample after confirmation with rarefaction curves (Supplemental Figure S1). Alpha-diversity indexes (Chao1, Simpson diversity index, Shannon diversity index, Pielou’s index, and good coverage index) were calculated in QIIME2. To detect significant differences in alpha-diversity metrics, a Kruskal-Wallis tests with Hodges-Lehmann estimate was performed (p<0.05). For beta-diversity, to test statistical differences between locations within each condition (“PWN-infected” and “Non-infected”) on the wood dataset and between locations on the nematode dataset, a one-way PERMANOVA was carried out. For further analysis, filtered ASV tables in QIIME2 were exported to R version 4.2.1 [48] and analyzed with “Phyloseq” and “MicrobiotaProcess” packages [49-50]. To describe each dataset, a compositional and structural analysis was performed: Venn diagrams were created to display shared and unique ASVs among conditions and locations; the relative abundance of the top 20 orders was plotted; and a principal coordinate analysis (PCoA) using Bray-Curtis dissimilarity matrix was created to compare fungal assemblages in a multidimensional space. A SIMPER analysis (100% cut-off percentage) based on Bray-Curtis dissimilarity was also conducted using PRIMER v6 software [51] to infer the relative contribution of each taxon to the (dis)similarities between conditions.

### Detection of *Bursaphelenchus xylophilus* in *Pinus pinaster* samples using real-time PCR

Total DNA samples used for metagenomic analysis were also used for the detection of PWN by real-time PCR. Gene-specific primers for *B. xylophilus* detection [52] and the universal 18S primers [53] were used in qPCR reactions (15 μl) containing 7.5 μl of NZYSupreme qPCR Probe (NZYTech, Lisboa, Portugal), 0.09 μl each specific primer (50 μM), 0.03 μl specific probe (50 μM), and 10 ng first-strand cDNA. The real-time PCR program included an initial denaturation for 10 min at 95 °C followed by 30 cycles of denaturation at 95 °C for 15 s, annealing and extension for 60 s at 60 °C. Two technical replicates were analyzed for each biological replicate. A positive amplification control (PAC, *B. xylophilus* DNA) and a negative amplification control (NAC, PCR-grade water) were included in all assays.

## Data Availability

The datasets presented in this study can be found in NBCI Bioproject database (PRJNA975140) under accession numbers SAMN31682492, SAMN38143581-SAMN38143583.

## Acknowledgments

The authors would like to acknowledge the researchers Luis Bonifácio (INIAV) and Soraia Vieira (MARE-UE) for their contributions in, respectively, the selection and collection of *Pinus pinaster* samples in all three sites, and in the revision of the statistical analysis of high-throughput sequencing data.

## Author contributions statement

CSVL and MLI: conceptualization. CSVL and ARV: research and data analysis. CSLV and ME: writing – original draft preparation. CSLV, ME, ARV, MLI: writing – review and editing. ME and MLI: resources. All authors contributed to the article and approved the manuscript.

