## Supplemental data for "Mycobiome of *Pinus pinaster* trees naturally infected by the pinewood nematode *Bursaphelenchus xylophilus*"

**(a)**

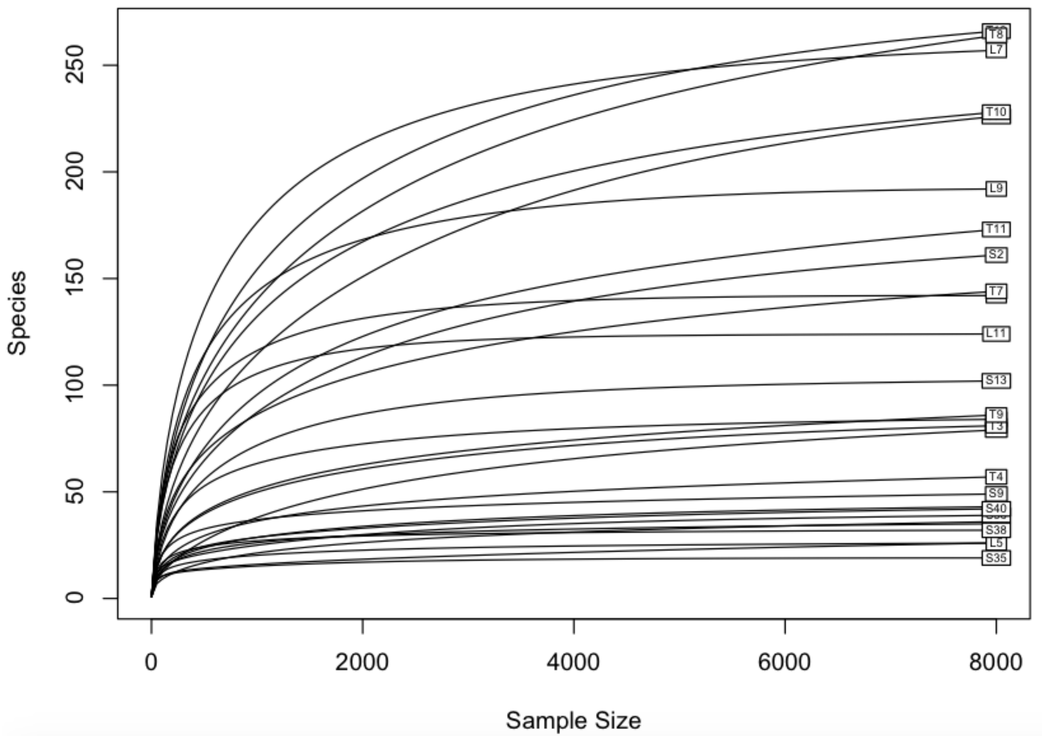


**(b)**

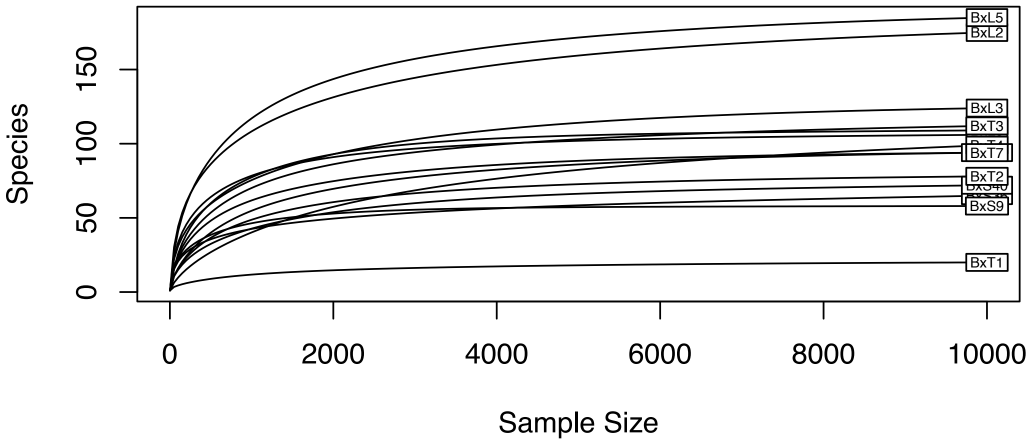


**Supplemental Figure S1. Rarefaction curves of the ASVs at 99% similarity for each sample** (a, represents non-infected and PWN-infected Pinus pinaster trees; b, represents PWN samples).

**Supplemental Table S1. Alpha-diversity estimates for each sample in both datasets.** Samples highlighted in grey correspond to *Pinus pinaster* trees infected with *Bursaphelenchus xylophilus*.

| | | Diversityestimates | | | | | |
|---|---|---|---|---|---|---|---|
| **Location** | **Samples** | **ObservedASVs[a]** | **Chao1** | **Simpson'sindex (1-D)** | **Shannon's index(H)** | **Pielou index** | **Good's Coverage** |
| Companhia das Lezírias | L2 | 31 | 31 | 0.79 | 1.99 | 0.58 | 1.00 |
| | L3 | 22 | 22 | 0.67 | 1.59 | 0.48 | 1.00 |
| | L4 | 42 | 45 | 0.79 | 1.97 | 0.51 | 1.00 |
| | L5 | 25 | 25 | 0.72 | 1.73 | 0.54 | 1.00 |
| | L6 | 32 | 32 | 0.53 | 1.28 | 0.34 | 1.00 |
| | L7 | 249 | 256 | 0.97 | 4.52 | 0.81 | 1.00 |
| | L8 | 142 | 142 | 0.97 | 4.14 | 0.84 | 1.00 |
| | L9 | 188 | 197 | 0.95 | 4.15 | 0.80 | 1.00 |
| | L11 | 124 | 126 | 0.96 | 3.91 | 0.81 | 1.00 |
| | BxL2 | 179 | 188 | 0.97 | 4.02 | 0.77 | 1.00 |
| | BxL3 | 118 | 125 | 0.94 | 3.34 | 0.69 | 1.00 |
| | BxL4 | 104 | 104 | 0.94 | 3.43 | 0.73 | 1.00 |
| | BxL5 | 190 | 196 | 0.94 | 3.71 | 0.70 | 1.00 |
| | BxL6 | 110 | 111 | 0.63 | 2.21 | 0.46 | 1.00 |
| Seia | S9 | 45 | 45 | 0.87 | 2.59 | 0.68 | 1.00 |
| | S35 | 18 | 18 | 0.77 | 1.81 | 0.63 | 1.00 |
| | S36 | 37 | 38 | 0.88 | 2.42 | 0.65 | 1.00 |
| | S38 | 32 | 32 | 0.86 | 2.40 | 0.69 | 1.00 |
| | S40 | 41 | 44 | 0.79 | 2.09 | 0.58 | 1.00 |
| | S1 | 74 | 89 | 0.69 | 1.65 | 0.37 | 1.00 |
| | S2 | 148 | 158 | 0.85 | 2.96 | 0.59 | 1.00 |
| | S13 | 99 | 100 | 0.71 | 2.32 | 0.51 | 1.00 |
| | S32 | 206 | 215 | 0.93 | 3.51 | 0.65 | 0.99 |
| | BxS9 | 61 | 61 | 0.86 | 2.63 | 0.64 | 1.00 |
| | BxS36 | 67 | 68 | 0.88 | 2.64 | 0.62 | 1.00 |
| | BxS38 | 91 | 94 | 0.71 | 2.25 | 0.50 | 1.00 |
| | BxS40 | 71 | 72 | 0.77 | 2.01 | 0.47 | 1.00 |
| Tróia | T1 | 82 | 82 | 0.88 | 2.88 | 0.65 | 1.00 |
| | T3 | 74 | 80 | 0.68 | 2.11 | 0.49 | 1.00 |
| | T4 | 55 | 59 | 0.79 | 2.12 | 0.53 | 1.00 |
| | T7 | 134 | 162 | 0.92 | 3.31 | 0.67 | 1.00 |
| | T8 | 237 | 251 | 0.96 | 4.06 | 0.74 | 0.99 |
| | T9 | 80 | 88 | 0.66 | 2.08 | 0.47 | 1.00 |
| | T10 | 217 | 229 | 0.95 | 3.92 | 0.71 | 1.00 |
| | T11 | 162 | 176 | 0.92 | 3.29 | 0.64 | 1.00 |
| | T12 | 256 | 269 | 0.96 | 4.19 | 0.76 | 1.00 |
| | BxT1 | 21 | 22 | 0.36 | 0.67 | 0.22 | 1.00 |
| | BxT2 | 78 | 79 | 0.77 | 2.03 | 0.47 | 1.00 |
| | BxT3 | 112 | 120 | 0.83 | 2.67 | 0.56 | 1.00 |
| | BxT4 | 108 | 114 | 0.34 | 0.97 | 0.21 | 1.00 |
| | BxT7 | 97 | 103 | 0.65 | 1.84 | 0.39 | 1.00 |

Chao1: Chao's species richness estimator
Shannon: Shannon diversity index
[a]Specieslevel:99%similarity threshold to define average sequence variants (ASVs)

**Supplemental Table S2. SIMPER analysis of similarity (Sim) and dissimilarity (diss) for *Pinus pinaster* dataset, considering as group factor disease condition (90% cut-off).** Only the top 3 species (ASV, average sequence variants) are presented.

| SIMPER Analysis | Condition | Av. Sim | Species | Av. Abund | Av. Sim | Sim/SD | Contrib% | Cum% |
|---|---|---|---|---|---|---|---|---|
| Similarity | PWN-infected | 20.06 | ASV0014 | 21.72 | 2.64 | 0.77 | 13.18 | 13.18 |
| | | | ASV0153 | 12.86 | 2.39 | 1.65 | 11.94 | 25.11 |
| | | | ASV0935 | 11.99 | 2.20 | 1.41 | 10.98 | 36.10 |
| | Non-infected | 18.96 | ASV0127 | 8.61 | 0.92 | 2.70 | 4.87 | 4.87 |
| | | | ASV0626 | 12.90 | 0.67 | 0.84 | 3.51 | 8.38 |
| | | | ASV0310 | 9.75 | 0.65 | 0.88 | 3.44 | 11.82 |
| **SIMPER Analysis** | **Groups** | **Av. Diss** | **Species** | **Av. Abund** | **Av. Diss.** | **Diss./SD** | **Contrib%** | **Cum%** |
| Dissimilarity | PWN-infected vs Non-infected | 94.59 | ASV0014 | 21.72 | 1.82 | 1.10 | 1.92 | 1.92 |
| | | | ASV0131 | 19.27 | 1.79 | 0.87 | 1.90 | 3.82 |
| | | | ASV0502 | 14.35 | 1.50 | 0.71 | 1.59 | 5.40 |

**Supplemental Table S4. SIMPER analysis of similarity (Sim) and dissimilarity (diss) for *Bursaphelenchus xylophilus* dataset, considering as group factor the locations (90% cut-off).** Only the top 3 species (ASV, average sequence variants) are presented.

| SIMPER Analysis | Location | Av. Sim | Species | Av. Abund | Av. Sim | Sim/SD | Contrib% | Cum% |
|---|---|---|---|---|---|---|---|---|
| Similarity | C. Lezírias(L) | 36.76 | ASV0607 | 19.49 | 1.85 | 2.81 | 5.02 | 5.02 |
| | | | ASV0447 | 13.27 | 1.48 | 6.86 | 4.02 | 9.05 |
| | | | ASV0609 | 15.27 | 1.41 | 2.64 | 3.82 | 12.87 |
| | Seia(S) | 43.51 | ASV0543 | 53.88 | 8.64 | 2.80 | 19.85 | 19.85 |
| | | | ASV0464 | 23.04 | 3.46 | 5.47 | 7.94 | 27.79 |
| | | | ASV0606 | 21.39 | 3.12 | 3.26 | 7.18 | 34.97 |
| | Tróia(T) | 29.96 | ASV0140 | 50.26 | 7.98 | 0.93 | 26.64 | 26.64 |
| | | | ASV0066 | 42.68 | 4.90 | 1.00 | 16.36 | 43.00 |
| | | | ASV0458 | 27.96 | 4.49 | 1.00 | 15.00 | 58.00 |
| **SIMPER Analysis** | **Location Groups** | **Av. Diss** | **Species** | **Av. Abund** | **Av. Diss.** | **Diss./SD** | **Contrib%** | **Cum%** |
| Dissimilarity | L&S | 71.08 | ASV0543 | 11.52 | 53.88 | 3.34 | 2.21 | 4.70 |
| | | | ASV0606 | 3.06 | 21.39 | 1.44 | 1.90 | 2.03 |
| | | | ASV0139 | 15.66 | 0.00 | 1.41 | 0.49 | 1.99 |
| | L&T | 86.71 | ASV0140 | 2.06 | 50.26 | 4.35 | 1.28 | 5.02 |
| | | | ASV0066 | 0.00 | 42.68 | 3.35 | 1.44 | 3.87 |
| | | | ASV0458 | 0.55 | 27.96 | 2.42 | 1.47 | 2.79 |
| | S&T | 89.46 | ASV0543 | 53.88 | 0.00 | 5.94 | 3.01 | 6.64 |
| | | | ASV0140 | 3.29 | 50.26 | 5.72 | 1.22 | 6.39 |
| | | | ASV0066 | 0.00 | 42.68 | 4.29 | 1.47 | 4.80 |
